# B cells engineered to express an anti-HIV antibody allow memory retention, class switch recombination and clonal selection in mice

**DOI:** 10.1101/2020.02.28.970822

**Authors:** Alessio D. Nahmad, Yuval Raviv, Miriam Horovitz-Fried, Ilan Sofer, Tal Akriv, Daniel Nataf, Iris Dotan, David Burstein, Yariv Wine, Itai Benhar, Adi Barzel

**Affiliations:** The School of Neurobiology, Biochemistry and Biophysics, The George S. Wise Faculty of Life Sciences, Tel Aviv University, Tel Aviv 69978, Israel; The School of Molecular Cell Biology and Biotechnology, The George S. Wise Faculty of Life Sciences, Tel Aviv University, Tel Aviv 69978, Israel

## Abstract

HIV viremia can be controlled by chronic antiretroviral therapy. As a potentially single-shot alternative, B cells engineered by CRISPR/Cas9 to express anti-HIV broadly neutralizing antibodies (bNAbs) were shown capable of secreting high antibody titers. Here, we demonstrate that, upon immunization of mice, adoptively transferred engineered B cells home to germinal centers (GC) where they predominate over the endogenous response and differentiate into memory and plasma cells while undergoing class switch recombination (CSR). Immunization with a higher affinity antigen increases accumulation in GCs and CSR rates. Boost immunization increases rates of engineered B cells in GCs and antibody secretion, indicating memory retention. Finally, antibody sequences of engineered B cells in the spleen show patterns of clonal selection. B cells may thus be engineered as a living and evolving drug.

## Main Text

Chronic antiretroviral therapy does not eradicate HIV infection. Broadly neutralizing antibodies (bNAbs) can suppress viremia^1^, but they may have to be chronically administered at a higher cost. Alternatively, bNAbs may be constitutively expressed from muscle^2,3^, but anti-drug antibodies (ADA) are often developed ^4^, possibly due to improper glycosylation. In addition, antibodies expressed from muscle undergo neither class switch recombination (CSR) nor affinity maturation, which may be necessary for long term control over diverse and continuously evolving HIV infections. These challenges may be overcome by B cell engineering. A therapeutically relevant protocol was first accomplished by lentiviral transduction of human CD34^+^ cells followed by *in vitro* differentiation^5^. Lentiviral transduction of mature B cells allowed the developmentally regulated expression of the membranal and secreted antibody isoforms^6^. More recently, efficient CRISPR/Cas9-mediated integration of antibody genes was demonstrated into the Ig loci of primary human B cells^7^. Integration of an antibody’s variable heavy chain into the immunoglobulin heavy (IgH) locus further allows somatic hyper-mutation (SHM) and class switch recombination *in vitro* when the endogenous constant segments are utilized using appropriate splicing signals^8^. In immunocompetent mice, adoptive transfer of B cells, engineered to express HIV-bNAbs, facilitated the production of HIV-neutralizing antibody titers^9^. Integration of single-chain anti-RSV antibodies into the IgH locus further allowed protection from infection^10^. However, in all previous *in vivo* studies neither immunological memory nor clonal selection could be detected, significantly hindering clinical application of the technology against the highly diverse and rapidly evolving HIV.

To overcome these limitations, we combine Toll-like receptor (TLR)-mediated *ex vivo* activation with *in vivo* prime-boost immunization. Uniquely, we demonstrate immunological memory and clonal selection that may contribute to addressing viral variability between patients and to counteracting viral escape. In particular, we use CRISPR/Cas9 and recombinant adeno associated viral vectors (rAAV) to target the integration of the 3BNC117^11^ HIV-bNAb under an Enhancer Dependent (ED) Ig promoter^12^ into the J-C intron of the IgH locus (Fig. 1A). We chose 3BNC117, a potent CD4-mimic HIV-bNAb, because, in combination with the 10-1074 bNAb, it was recently shown to induce viral suppression in viremic individuals^13^ and in individuals undergoing treatment interruption^14^. Our bi*-*cistronic bNAb cassette encodes the full light chain and the variable segment of the heavy chain (VH) of 3BNC117 separated by a furin cleavage site and a 2A-peptide for ribosomal skipping. The VH is followed by a splice donor sequence to allow fusion to constant segments and initial expression of the bNAb as a membranal B cell receptor (BCR). Our design may allow disruption of the endogenous IgH chain while facilitating antigen-induced activation of engineered B cells upon immunization, leading to differentiation into memory and plasma cells, as well as to CSR, SHM and affinity maturation (Fig. 1A).

**Fig. 1.**
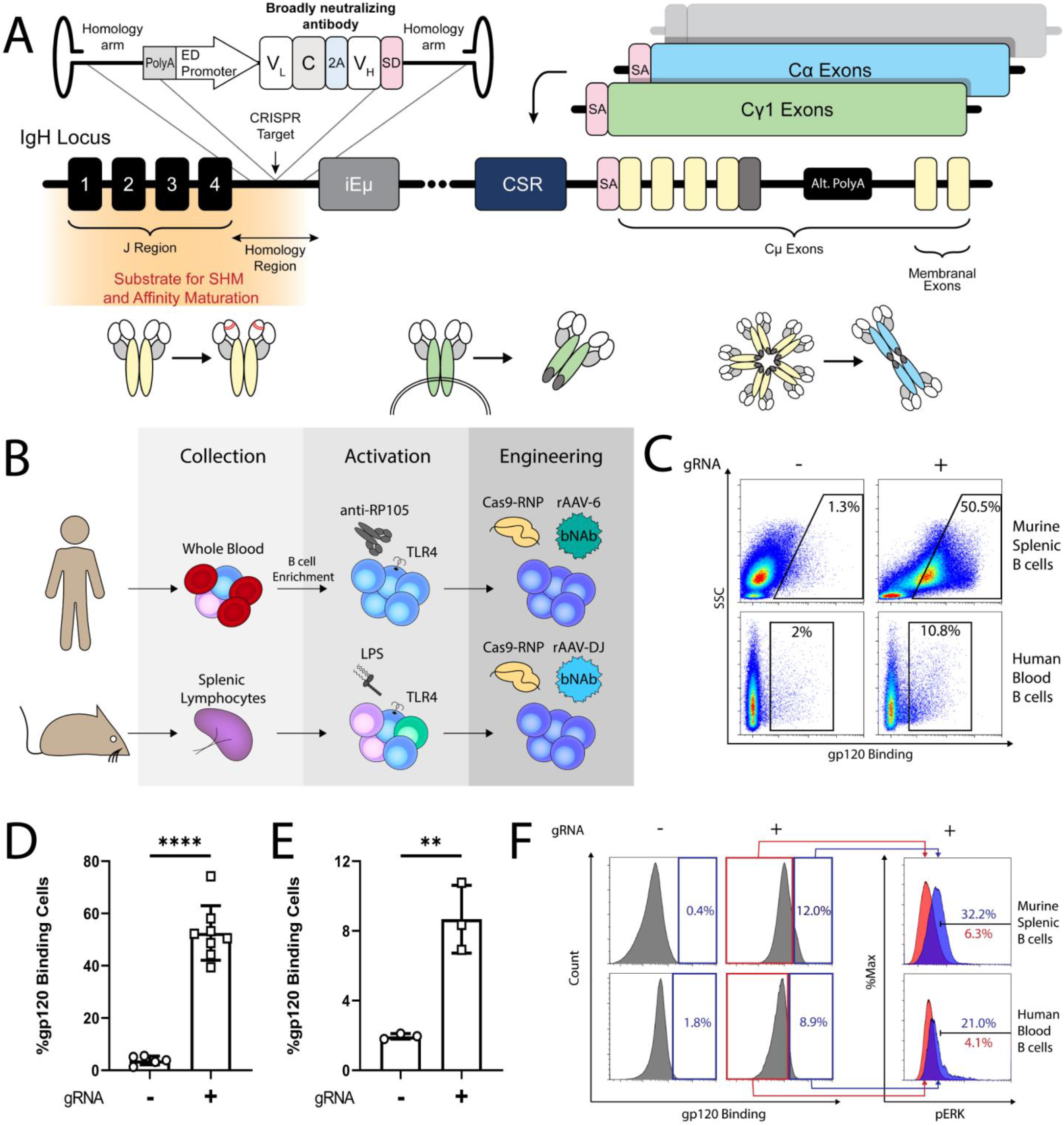
Engineering B cells to express an anti-HIV bNAb **(A)** Targeting scheme. An rAAV-delivered cassette is targeted to the J-C intron of the IgH locus using CRISPR/Cas9. The bicistronic cassette encodes the light and heavy chains of the 3BNC117 anti-HIV bNAb, under the control of an enhancer dependent (ED) promoter. Splicing with endogenous constant segments allows the expression of a BCR and differentiation into memory B cells and Ig secreting plasma cells upon subsequent antigen-induced activation and alternative polyadenylation (Alt. PolyA). Targeting the J-C intron upstream of the intronic enhancer (iEμ) and switch region further facilitates CSR and SHM **(B)** Activation and engineering scheme. Human B cells are collected from blood samples, activated using an anti-RP105 (TLR4 homologue) antibody, electroporated by CRISPR/Cas9 RNP and transduced using rAAV-6. Splenic B cells are activated using the TLR4 agonist LPS electroporated by CRISPR/Cas9 RNP and transduced using rAAV-DJ. **(C)** Flow cytometry plots measuring binding of the HIV gp120 antigen by the 3BNC117 BCR following activation and engineering of primary cells. Cells transduced with the donor rAAV and without gRNA serve as a negative control, gating on live, singlets. **(D and E)** Quantification of C for murine **(D)** and human **(E)** primary cells. Each dot represents an independent assay, **** = pv<0.0001, ** = pv<0.001; t-test. **(F)** Flow cytometric analysis of ERK phosphorylation in primary murine or human B cells engineered with 3BNC117 and *in-vitro* activated with the gp120 antigen of the YU2.DG HIV strain, gating on singlets.

First, CRISPR/Cas9 gRNAs were designed to promote cleavage of the murine and the human IgH J-C introns. We chose the specific target positions to allow donor-vector design with sufficiently long homology arms that contain neither the potentially oncogenic intronic-enhancer nor sequences corresponding to genomic segments that may be deleted during VDJ recombination. CRISPR/Cas9 cleavage produced up to 33% and 21% InDels, by the T7E1 assay, at the IgH locus of an immortalized pro B cell line (ImProB)^15^ and the Ramos human B cell line, respectively (Fig. S1A). Different gRNAs were used to disrupt the respective endogenous kappa light chains (IgK) in order to avoid chain mispairing. The IgK gRNAs target the splice acceptor junctions of the IgK constant segments, to avoid targeting the IgK chain of the transgenic bNAb. Following CRISPR/Cas9 cleavage, IgK expression was ablated in up to 19% and 16% in the murine and human cell lines, respectively (Fig. S1B). Next, we combined the CRISPR/Cas9 ribonucleoprotein (RNP) electroporation with rAAV transduction. We first used an rAAV encoding a GFP-cassette to validate that the activity of our Ig promoter variant is dependent upon on-target integration next to the Intronic enhancer. Indeed, high and stable GFP expression was demonstrated in the ImProB cell line, but only when the appropriate gRNA was co-delivered (Fig. S2). When using the bNAb cassette (Fig. 1A), the combined CRISPR/Cas9 electroporation and rAAV transduction resulted in 3BNC117 BCR expression in 9% and 8% of cells in the murine and human lines, respectively (Fig. S3A). mRNA analysis confirmed splicing of the integrated 3BNC117 with the expected constant segment in each cell line (Fig. S3B).

Efficient engineering of primary B cells may require activation through either the CD40 pathway^10,8^ or the TLR pathway^9^. CD40 ligation can produce germinal center (GC) -like B cells and antibody secreting plasma cells *ex vivo* ^16^. However, B cells activated *ex vivo* through CD40 ligation express high levels of CD80, which are associated with a reduced propensity for further activation *in vivo* upon immunization^17,18^. In particular, these cells have a diminished capacity to home to GCs, establish immunological memory and undergo SHM and affinity maturation^10,18^. In contrast, TLR-pathway mediated activation of donor B cells, coming from a transgenic mouse, was recently shown to allow homing to recipient’s GCs upon immunization^9^. We therefore activated murine splenic B cells using the TLR4 agonist lipopolysaccharide (LPS) and activated human blood B cells using an antibody against the TLR4 homologue RP105 (Fig. 1B). CRISPR*/*Cas9 RNP electroporation led to 30% and 29% InDel formation at the mouse and human IgH loci, respectively (Fig S4A), and led to IgK ablation in 49% and 45% of cells, in the respective species (Fig. S4B) Following a subsequent rAAV transduction, up to 74% and 11% of cells expressed the 3BNC117 BCR amongst the activated murine and human cells, respectively (Fig. 1C-E). Correct integration was validated by Sanger sequencing (Fig. S4C). The distributions of phenotypes and isotypes were similar among 3BNC117 expressing and non-expressing cells (Fig. S4D and E). Importantly, as we activated the cells through the TLR rather than CD40 pathway, the engineered cells express low levels of CD80 (S4F), implying increased propensity for antigen-induced activation. 3BNC117 secretion was minor but detectable (Fig. S4G). In addition, high viability was retained following activation, electroporation and transduction of both mouse and human cells (Fig. S4H). Engineered B cells of both species could be further activated *ex vivo* by incubation with the HIV gp120 antigen, leading to ERK phosphorylation (Fig. 1F). We validated that the promoter we used is active in the primary cells only upon CRISPR-dependent integration next to the intronic enhancer (Fig. S5A). As a possible additional precaution, we demonstrated efficient B cell engineering with a promoter-less 3BNC117 cassette, allowing expression only upon integration into the IgH locus followed by splicing with the endogenous transcript (Fig. S5B-C, top). Similarly, the risk of mispairings between transgenic and endogenous chains may be reduced by coding 3BNC117 as a single chain antibody^10^ (Fig. S5B-C, bottom, S5D). While the promoter-less design was associated with a reduced engineering rate, the single-chain design allowed for a high rate of engineering.

In order to assess functionality *in vivo*, we first engineered activated splenic B cells of CD45.1 C57BL/6 mice to express 3BNC117, and then we adoptively transferred the cells into an otherwise syngeneic CD45.2 mice. Each recipient mouse received 1.5-2.2M donor cells, such that the number of 3BNC117 expressing cells transferred was set at 112,500. Different groups of mice subsequently received only prime or both prime and boost immunizations and were terminally bled to assess serum antibody concentration or sacrificed to analyze the splenic B cell population (Fig. 2A). Different groups were immunized with the gp120 antigen of the YU2.DG HIV strain, efficiently neutralized by 3BNC117, or by the gp120 antigen of the THRO4156.18 HIV strain, which is poorly neutralized by 3BNC117^19^. 3BNC117 has a much higher affinity to the YU2.DG antigen (Fig. S6), but the antigens are otherwise comparable, both belonging to the clade B HIV-1 strains. While less than 10% of the transferred cells were gp120 binders, following immunization with either antigen, the vast majority of CD45.1 expressing cells in the GCs bound gp120 (Fig 2B-C), a strong indication for antigen-induced homing to GCs. Furthermore, while gp120 is immunogenic to mice, irrespective of B cell engineering (Fig. S7), 8 days following prime immunization with either antigen, relevant splenic GCs were monopolized by donor cells, with more than 90% of gp120 binding cells expressing CD45.1 (Fig 2D-E). Therefore, in patients, activation of engineered B cells encoding neutralizing antibodies may come at the expense of the endogenous response, which is often non-neutralizing^20,21^.

**Fig. 2.**
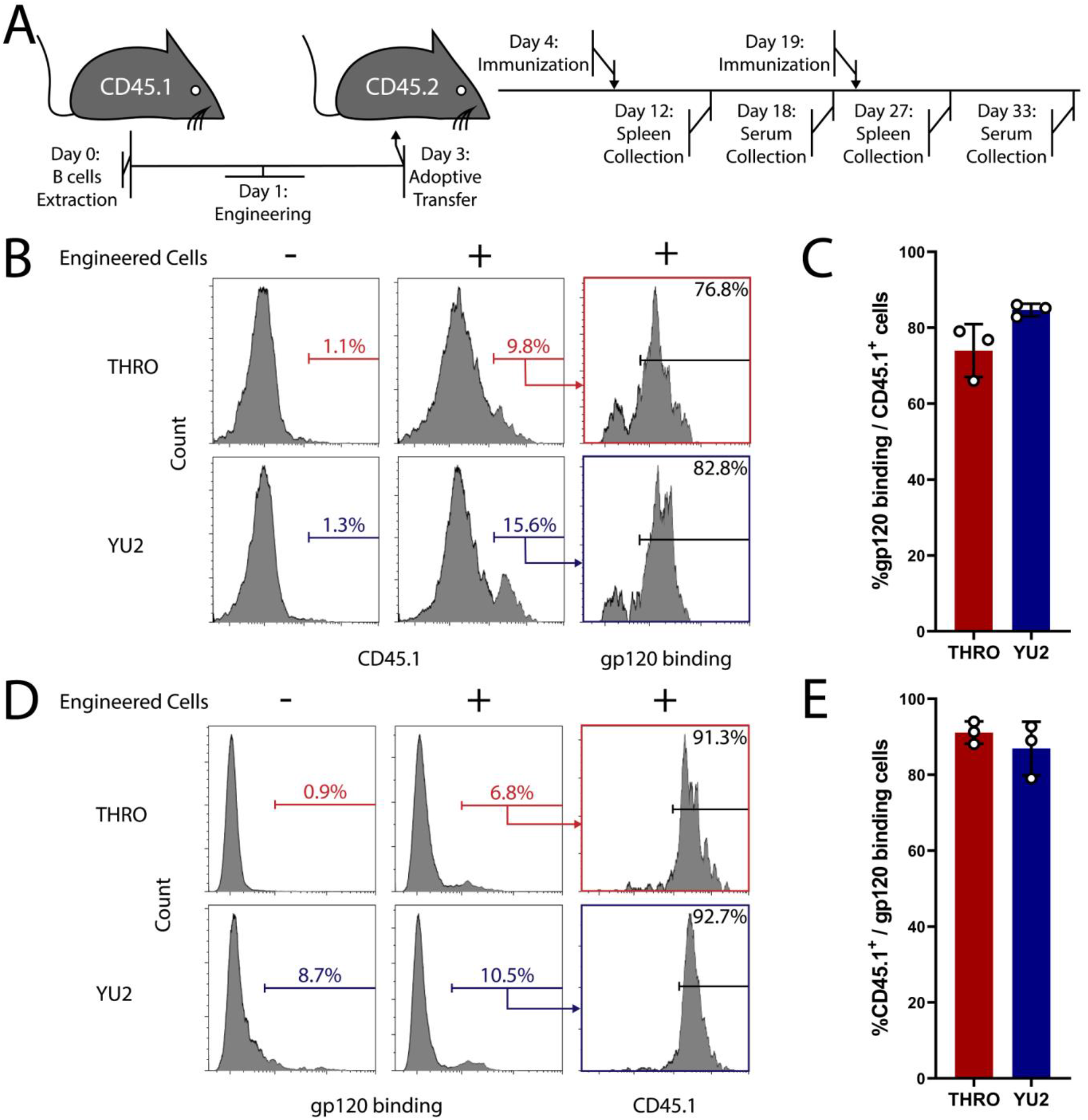
Adoptively transferred engineered B cells can undergo antigen-induced activation *in-vivo* **(A)** Experimental scheme of the *in vivo* assays. Splenic B cells, from C57BL/6 CD45.1 mice, were engineered as in Fig.1 and infused to otherwise syngeneic CD45.2 recipient mice. Different mice groups were immunized on the following day with gp120 antigens from either the THRO4156.18 (THRO) or the YU2.DG (YU2) HIV strains. When boosted by an additional injection, the mice received the same antigen as in the prime injection. Different mice groups were sacrificed 8 days following injections for spleen collection or terminally bled 14 days after injection for serum collection. **(B)** Representative analysis by flow cytometry of the accumulation of engineered cells in the GCs of mice immunized with either the YU2.DG or the THRO4156.18 gp120 antigens, 8 days following a prime antigen injection. Gating on live, singlets, B220^+^, GL-7^+^. **(C)** Quantification of B. Each dot represents a different mouse. Error bars represent SD. **(D)** Representative analysis by flow cytometry of CD45.1 expression among gp120 binding GC cells. Pre-gating on singlets, live, B220^+^, GL-7^+^ **(E)** Quantification of D. Each dot represents a different mouse. Error bars represent SD. For gating strategy see Fig.S15.

14 days post-immunizations with the YU2.DG gp120 antigen, we found higher serum concentrations of the 3BNC117 Ab compared to the concentrations following immunizations with the THRO4156.18 antigen (Fig. 3A). However, importantly, a log increase in serum Ab concentrations was measured 14 days following boost immunizations, reaching nearly 1μg/ml using either of the antigens (Fig. 3A,S8). Such high 3BNC117 serum concentrations were previously demonstrated to allow broad HIV neutralization^22^. The rate of engineered cells in the GCs has also significantly increased 8 days following boost immunization with either of the antigens (Fig. 3B-C). Compared to mice receiving both prime and boost immunizations, rates of engineered cells in the GCs were lower in mice analyzed at a late time point after receiving only an early prime immunization and in mice receiving only a late prime immunization (Fig. S9A-B. Therefore, importantly, the boost effect can be strictly attributed to the retention of immunological memory. Indeed, immunophenotyping of the donor cells in the spleen revealed that the engineered cells were differentiating into both CD138^+^ Ig secreting cells and CD38^+^ (which include memory B cells) (Fig. 3D-E,S9C). Interestingly, immunization with the YU2.DG gp120 antigen induced higher rates of differentiation into CD38^+^ B cell compared to immunization with the THRO4156.18 antigen. In addition, the rate of splenic CD38^+^ B cells was increased while the rate of plasma cells was decreased following boost immunization with either of the antigens, in concordance with natural murine boost responses^23^.

**Fig. 3.**
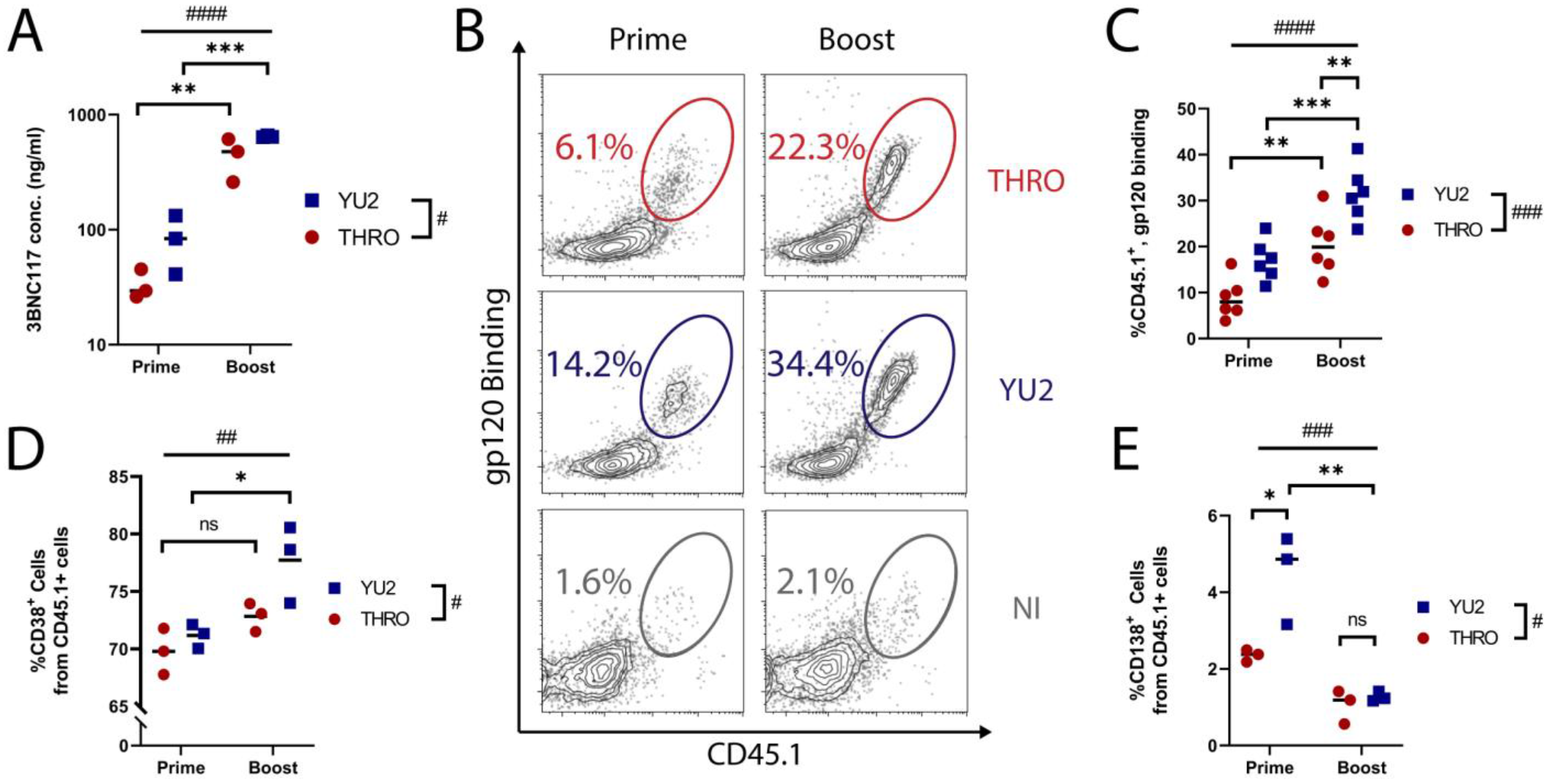
Adoptively transferred engineered B cells allow memory retention upon immunization **(A)** ELISA of sera collected 14 days following either prime or boost immunization, quantified using an anti-idiotypic antibody to 3BNC117. # # # # = pv<0.0001, # = pv<0.05 for two-way ANOVA and ** = pv<0.01, *** = pv<0.001 for Tukey’s multiple comparison **(B)** Analysis by flow cytometry of CD45.1 expression and gp120 binding in the GCs of mice following prime or boost immunizations, gating on live, singlets, B220^+^, GL-7^+^. NI: Non-immunized; recipient mice received engineered cells as in (Fig.2A) but were not immunized **(C)** Quantitation of B. # # # # = pv<0.0001, # # # = pv<0.001 for two-way ANOVA and *** = pv<0.001 and ** = pv<0.01 for Tukey’s multiple comparison. **(D-E)** Analysis by flow cytometry of CD38 or CD138 expression among donor derived cells in the spleens of recipient mice after prime or boost immunizations by the gp120 antigens from either the THRO4156.18 (THRO, Red) or the YU2.DG (YU2, Blue) HIV strains, gated on live, singlets, CD45.1^+^. # # # = pv<0.001, # # = pv<0.01, # = pv<0.05 for two-way ANOVA and ** =pv<0.01, * = pv<0.05 for Tukey’s multiple comparison. For gating strategy see Fig.S15.

CSR may be necessary to ensure both humoral and mucosal protection from HIV surge. Indeed, IgG_1_, IgG_2_ and IgA isotypes of the 3BNC117 bNAb were found in the sera of treated mice in addition to the IgM isotype (Fig. 4A, S10A-C). Class switched 3BNC117 antibodies were more prevalent in sera when the YU2.DG gp120 antigen was used for immunization, and engineered cells expressing the IgA isotype were found in the GCs of treated mice only upon prime immunization by the YU2.DG gp120 antigen (Fig 4B). As CSR often precedes GC homing^24^, this trend is in agreement with the higher rates of GC B cells in mice immunized by the YU2.DG antigen. Notably, rates of IgA expression among donor cells in the GCs, after immunizations with YU2.DG, were higher than the pre-implantation rates, implying antigen-induced *in vivo* CSR (Fig. 4B,S10D).

**Fig. 4.**
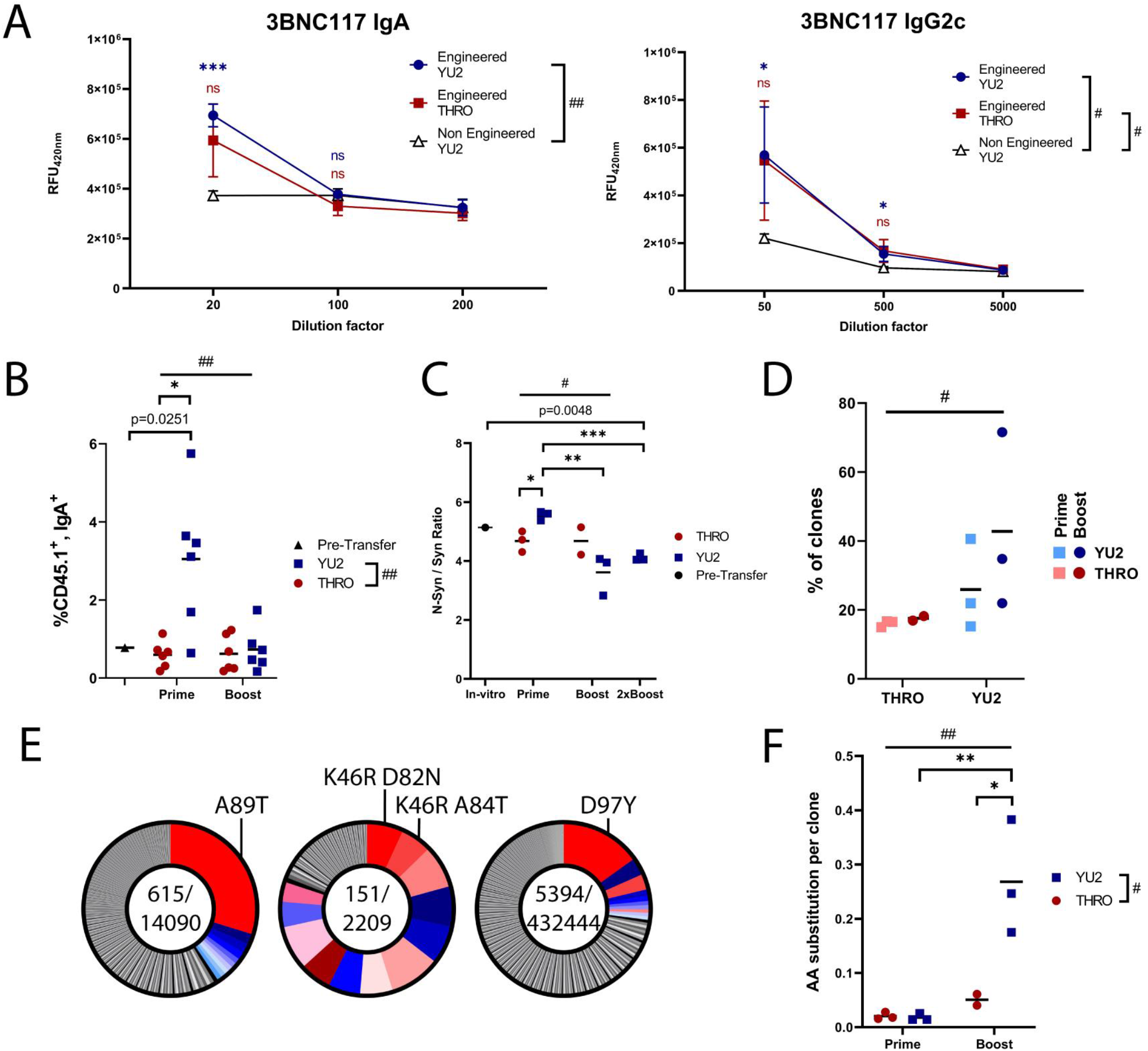
Adoptively transferred engineered B cells can undergo CSR and clonal expansion upon immunization **(A)** Isotype specific anti-idiotypic ELISA measuring 3BNC117 isotypes in mice sera collected after boost immunizations. # = pv<0.05, ## = pv<0.01 for Dunnett’s multiple comparisons and *** = pv<0.001 and * = pv<0.05 for t-test. Comparisons performed to sera of mice adoptively transferred with non-engineered B cells and boost immunized with the YU2.DG gp120 antigen. **(B)** Analysis by flow cytometry of IgA and CD45.1 expression among GC cells after prime and boost immunizations, gating on live, lymphocytes, GL-7^+^, B220^+^. ## = pv<0.01 for two-way ANOVA,* = pv<0.01 for t-test and indicated p value is for one-sample t-test **(C)** Ratio of non-synonymous to synonymous mutations in the different samples. # = pv<0.05 for two-way ANOVA between the prime and boost cohorts. * = pv<0.05, ** = pv<0.01, *** pv<0.001 for t-test and indicated p value is for one-sample t-test.**(D)** Quantitation of the clonal expansion by measuring polarity: the relative cumulative share of the 10 most abundant clones. # = pv<0.05; two-way ANOVA. **(E)** Pie charts of mice immunized with the YU2.DG gp120 antigen and having at least one clone representing more than >10% of the mutant repertoire. The most abundant clones in each mouse were colored. Shades of red indicate clones that were not found in the 10 most abundant clones of other mice (Fig.S13). Shades of blue indicate shared clones. Indicated clones are the K46R, A89T and D97Y. **(F)** Quantitation of amino acid (AA) substitutions per clone in mice prime- or boost-immunized with the YU2.DG or THRO4156.18 gp120 antigens. ## = pv < 0.01, # = pv<0.05 for two-way ANOVA and ** = pv<0.01, * = pv < 0.05 for Tukey’s multiple comparison.

Finally, in order to assess *in vivo* SHM and clonal expansion among engineered B cells, we used a synonymously re-coded 3BNC117 allele, enriched for sequence hotspots of activation-induced-cytidine-deaminase (AID, catalyzing SHM) (Fig. S11A-C). Accumulation in mouse GCs following immunizations was similar whether the adoptively transferred B cells were engineered to express 3BNC117-wt or the recoded variant: 3BNC117-opt (Fig. S11D). We harvested RNA from the spleens of mice receiving engineered cells and amplified the bNAb V_H_ sequence from the cDNA (Fig. S12A) for analysis by Illumina sequencing. We found strong evidence for clonal expansion. While some mutations were found to arise during AAV preparation and B cell engineering^25^, the distribution of mutations along the 3BNC117 sequence has shifted following adoptive transfer and immunizations in patterns implying *in-vivo* selection (Fig. S12B) SHM and/or *in vivo* selection are further implied by the decrease in the ratio of non-synonymous to synonymous mutations upon adoptive transfer followed by prime and boost immunizations with the YU2.DG gp120 antigen (Fig. 4C). Furthermore, clonal expansion following YU2.DG gp120 immunizations was evident by the marked increase in the relative share of the 10 most abundant clones (Fig. 4D, S13A), and the selection of clones was similar between mice from the same cohort (Fig. S13B-E). In particular, the three amino acid differences found in more than 10% of the variants, in their respective mice, were D97Y in the CDR3, A89T in a FR3/CDR3 border position and K46R in the CDR2 (Fig. 4E, S13F). Clonal expansions of the K46R and A89T substitutions are supported by their association with multiple additional substitutions (Fig. S14A-C) in their respective mice. None of these combinations of substitutions could be found in any other mouse, including in 5 mice adoptively transferred by the same pool of engineered B cells. This strongly indicates *de-novo* SHM. Clonal expansion of both the K46R and the A89T substitutions is further supported by the presence of the respective R and T amino acids in the sequence of the related VRC01 and NIH45-46 antibodies^26^ (Fig.S13F). Interestingly, immunizations with the higher affinity antigen, YU2.DG as well as boost immunizations triggered higher accumulation of amino acid substitutions (Fig. 4F). The limited clonal expansion following immunizations with the THRO4156.18 gp120 antigen may imply that multiple or rare substitutions in the 3BNC117 sequence are necessary in order to allow sufficient improvement in the affinity toward this antigen.

In conclusion, we have uniquely demonstrated that B cells engineered to express bNAbs can undergo antigen-induced activation in mice followed by memory retention, CSR, SHM and clonal expansion. In the human setting, bNAb SHM may facilitate affinity maturation to counteract HIV diversity and high mutation rate. Moreover, patient or donor B cells may be engineered to express two or more bNAbs^14^ targeting different viral epitopes, to further diminish the risk of escape. Both safety and efficiency may be increased by expressing the bNAb as a single chain and by using promoter-less constructs that more strictly prevent expression from off target integration^27^. The therapeutic potential of our approach may best be assessed in non-human primates with HIV-like infections. Finally, B cell engineering as a platform technology may be applied in the future to diverse persistent infections as well as to the treatment of congenital disorders, autoimmune diseases and cancer.

## Acknowledgments

We thank the Veterinary Service Center, Tel Aviv University for animal husbandry and the IDRFU and SICF units, Tel Aviv university for logistic support. We also thank Omri Wurtzel, Ziv Shulman, Natalia Freund, Steffen Jung, Adi Stern, Rina Rosin-Arbesfeld and Natalie Zelikson for reagents and feedback:

## Author contributions

A.D.N designed, performed and analyzed the study and also drafted and revised the manuscript; Y.R performed bioinformatics analysis; M.H-F. performed flow cytometry; T.A. and D.N. contributed to the experimental design; I.S. Designed constructs, I.D contributed to conceiving and supervising the study; D.B. and Y.W. Performed bioinformatic analysis, I.B. generated antibodies; A.B. Conceptualized and supervised the study.

## Materials and Methods

### crRNA and sgRNA for SpCas9 cleavage

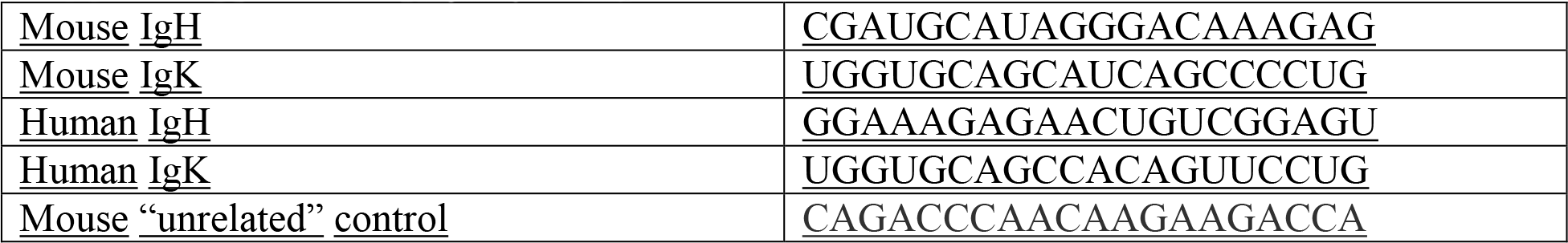

### Plasmid cloning

For the murine donor plasmid, two intronic sequences directly adjacent to the murine target gRNA (homology arms) were PCR amplified using primers ATTAATTAAGCGGCCGCGTAAGAATGGCCTCTCCAGGTCTT (forward) CTCCACTCCTCGAGTTTGTCCCTATGCATCG (reverse) and GGACAAACTCGAGGAGTGGAGTGGGGC (forward) ACGCGTGTACACTAGTCCAACTCAACATTGCTCAATTCATTTAAAAATATTTGAAACT

(reverse) from the ImProB cell line. Fragments were inserted by In-Fusion assembly into pAB270^1^ cut with NheI and SpeI. Finally, we inserted the sequence CTCGACTGTGCCTTCTAGTTGCCAGCCATCTGTTGTTTGCCCCTCCCCCGTGCCTTCCTTGACCCTGGAAGGTGCCACTCCCACTGTCCTTTCCTAATAAAATGAGGAAATTGCATCGCATTGTCTGAGTAGGTGTCATTCTATTCTGGGGGGTGGGGTGGGGCAGGACAGCAAGGGGGAGGATTGGGAAGACAATAGCAGGCATGCTGGGGATGCGGTGGGCTCTATGGGGTACCTTTTCATTCCTTCCTCTCCAGTTCTTCTCTAGATGGACTAGGTCCTTAACTAGCGAATTCGGATCCCTGTCTCATGAATATGCAAATCAGGTGAGTCCATGGTGGTAAATATAGGGATGTCGACACACCTCACAAACTTAAGATCTAGAATGGACATGAGGGTCCCTGCTCAGCTCCTGGGGCTCCTGCTGCTCTGGCTCTCTGGAGCCAGATGTGACATCCAGATGACCCAGTCTCCATCCTCCCTGTCTGCCTCTGTGGGAGATACCGTCACTATCACTTGCCAAGCAAACGGCTACTTAAATTGGTATCAACAGAGGCGAGGGAAAGCCCCAAAACTCCTGATCTACGATGGGTCCAAATTGGAAAGAGGGGTCCCATCAAGGTTTAGTGGAAGAAGATGGGGGCAAGAATATAATCTGACCATCAACAATCTGCAGCCCGAAGACATTGCAACATATTTTTGTCAAGTGTATGAGTTTGTCGTCCCTGGGACCAGACTGGATTTGAAACGGGCTGATGCTGCACCAACTGTATCCATCTTCCCACCATCCAGTGAGCAGTTAACATCTGGAGGTGCCTCAGTCGTGTGCTTCTTGAACAACTTCTACCCCAAAGACATCAATGTCAAGTGGAAGATTGATGGCAGTGAACGACAAAATGGCGTCCTGAACAGTTGGACTGATCAGGACAGCAAAGACAGCACCTACAGCATGAGCAGCACCCTCACGTTGACCAAGGACGAGTATGAACGACATAACAGCTATACCTGTGAGGCCACTCACAAGACATCAACTTCACCCATTGTCAAGAGCTTCAACAGGAATGAGTGTCGCGCGAAACGCGGAAGCGGAGCTACTAACTTCAGCCTGCTGAAGCAGGCTGGAGACGTGGAGGAGAACCCTGGACCTATGGACTGGACCTGGAGGATCCTCTTCTTGGTGGCAGCAGCCACAGGAGCCCACTCCCAGGTCCAATTGTTACAGTCTGGGGCAGCGGTGACGAAGCCCGGGGCCTCAGTGAGAGTCTCCTGCGAGGCTTCTGGATACAACATTCGTGACTACTTTATTCATTGGTGGCGACAAGCCCCAGGACAGGGCCTTCAGTGGGTGGGATGGATCAATCCTAAGACAGGACAGCCAAACAATCCTCGTCAATTTCAAGGTAGAGTCAGTCTGACTCGA CACGCGTCGTGGGACTTTGACACATTTTCCTTTTACATGGACCTGAAGGCACTAAGATCGGACGACACGGCCGTTTATTTCTGTGCGCGACAGCGCAGCGACTATTGGGATTTCGACGTCTGGGGCAGTGGAACCCAGGTCACTGTCTCGTCAGGTGAGTCCTCGAG for the 3BNC117-wt donor or: CTCGACTGTGCCTTCTAGTTGCCAGCCATCTGTTGTTTGCCCCTCCCCCGTGCCTTCCTTGACCCTGGAAGGTGCCACTCCCACTGTCCTTTCCTAATAAAATGAGGAAATTGCATCGCATTGTCTGAGTAGGTGTCATTCTATTCTGGGGGGTGGGGTGGGGCAGGACAGCAAGGGGGAGGATTGGGAAGACAATAGCAGGCATGCTGGGGATGCGGTGGGCTCTATGGGGTACCTTTTCATTCCTTCCTCTCCAGTTCTTCTCTAGATGGACTAGGTCCTTAACTAGCGAATTCGGATCCCTGTCTCATGAATATGCAAATCAGGTGAGTCCATGGTGGTAAATATAGGGATGTCGACACACCTCACAAACTTAAGATCTAGAATGGACATGAGGGTCCCTGCTCAGCTCCTGGGGCTCCTGCTGCTCTGGCTCTCTGGAGCCAGATGTGACATCCAGATGACACAGAGCCCTAGCAGCCTGTCTGCCAGCGTGGGAGACACCGTGACAATTACCTGCCAAGCTAACGGCTACCTGAACTGGTATCAGCAGAGAAGAGGCAAGGCCCCTAAGCTGCTGATCTACGACGGCAGCAAGCTGGAAAGAGGCGTGCCCTCTAGATTCAGCGGCAGAAGATGGGGCCAAGAGTACAACCTGACCATCAACAACCTGCAGCCTGAGGATATCGCCACATACTTCTGCCAAGTGTACGAGTTCGTGGTGCCCGGCACCAGACTGGACCTGAAGCGGGCTGATGCTGCACCAACTGTATCCATCTTCCCACCATCCAGTGAGCAGTTAACATCTGGAGGTGCCTCAGTCGTGTGCTTCTTGAACAACTTCTACCCCAAAGACATCAATGTCAAGTGGAAGATTGATGGCAGTGAACGACAAAATGGCGTCCTGAACAGTTGGACTGATCAGGACAGCAAAGACAGCACCTACAGCATGAGCAGCACCCTCACGTTGACCAAGGACGAGTATGAACGACATAACAGCTATACCTGTGAGGCCACTCACAAGACATCAACTTCACCCATTGTCAAGAGCTTCAACAGGAATGAGTGTCGCGCGAAACGCGGAAGCGGAGCTACTAACTTCAGCCTGCTGAAGCAGGCTGGAGACGTGGAGGAGAACCCTGGACCTATGGACTGGACCTGGAGGATCCTCTTCTTGGTGGCAGCAGCCACAGGAGCCCACTCCCAGGTTCAGCTGCTGCAATCTGGCGCCGCTGTGACAAAACCTGGCGCCTCTGTTAGAGTGTCCTGCGAGGCTAGCGGCTACAACATCAGAGACTACTTCATCCACTGGTGGCGGCAGGCTCCAGGACAGGGACTTCAATGGGTCGGCTGGATCAACCCCAAAACCGGGCAGCCTAACAACCCCAGACAGTTCCAGGGCAGAGTGTCCCTGACAAGACACGCCAGCTGGGACTTCGACACCTTCAGCTTCTACATGGACCTGAAGGCCCTGAGAAGCGACGATACAGCCGTGTACTTCTGTGCTCGGCAGCGGTCCGACTACTGGGACTTCGACGTGTGGGGCAGTGGAACCCAAGTTACTGTTTCGTCAGGTGAGTCCTCGAG for the 3BNC117-opt donor in the XhoI restriction site by Gibson Assembly (NEB).

For the human donor plasmid, the intronic sequences were amplified using primers CGCGATGCATTAATTAAGCGGCCGCGTAAGAATGGCCACTCTAGGGCC (forward) CCCACTCCCTCGAGGACAGTTCTCTTTCC (reverse) and AACTGTCCTCGAGGGAGTGGGTGAATCC (forward) GATATCACGCGTGTACACTAGTACAGCACTGTGCTAGTATTTCTTAGCT (reverse) and Gibson Assembled into pAB270 NheI-SpeI restricted. Sequence CCTCGACTGTGCCTTCTAGTTGCCAGCCATCTGTTGTTTGCCCCTCCCCCGTGCCTTCCTTGACCCTGGAAGGTGCCACTCCCACTGTCCTTTCCTAATAAAATGAGGAAATTGCATCGCATTGTCTGAGTAGGTGTCATTCTATTCTGGGGGGTGGGGTGGGGCAGGACAGCAAGGGGGAGGATTGGGAAGACAATAGCAGGCATGCTGGGGATGCGGTGGGCTCTATGGGGTACCTTTTCATTCCTTCCTCTCCAGTTCTTCTCTAGATGGACTAGGTCCTTAACTAGCGAATTCGGATCCCTGTCTCATGAATATGCAAATCAGGTGAGTCCATGGTGGTAAATATAGGGATGTCGACACACCTCACAAACTTAAGATCTAGAATGGACATGAGGGTCCCTGCTCAGCTCCTGGGGCTCCTGCTGCTCTGGCTCTCTGGAGCCAGATGTGACATCCAGATGACCCAGTCTCCATCCTCCCTGTCTGCCTCTGTGGGAGATACCGTCACTATCACTTGCCAAGCAAACGGCTACTTAAATTGGTATCAACAGAGGCGAGGGAAAGCCCCAAAACTCCTGATCTACGATGGGTCCAAATTGGAAAGAGGGGTCCCATCAAGGTTTAGTGGAAGAAGATGGGGGCAAGAATATAATCTGACCATCAACAATCTGCAGCCCGAAGACATTGCAACATATTTTTGTCAAGTGTATGAGTTTGTCGTCCCTGGGACCAGACTGGATTTGAAACGAACGGTGGCTGCACCATCTGTCTTCATCTTCCCGCCATCTGATGAGCAGTTGAA ATCTGGAACTGCCTCTGTTGTGTGCCTGCTGAATAACTTCTATCCCAGAGAGGCCAAAGTACAGTGGAAGGTGGATAACGCCCTCCAATCGGGCAACTCCCAGGAGAGTGTCACAGAGCAGGACAGCAAGGACAGCACCTACAGCCTCAGCAGCACCCTGACGCTGAGCAAAGCAGACTACGAGAAACACAAAGTCTACGCCTGCGAAGTCACCCATCAGGGCCTGAGCTCGCCCGTCACAAAGAGCTTCAACAGGGGAGAGTGTCGCGCGAAACGCGGAAGCGGAGCTACTAACTTCAGCCTGCTGAAGCAGGCTGGAGACGTGGAGGAGAACCCTGGACCTATGGACTGGACCTGGAGGATCCTCTTCTTGGTGGCAGCAGCCACAGGAGCCCACTCCCAGGTCCAATTGTTACAGTCTGGGGCAGCGGTGACGAAGCCCGGGGCCTCAGTGAGAGTCTCCTGCGAGGCTTCTGGATACAACATTCGTGACTACTTTATTCATTGGTGGCGACAAGCCCCAGGACAGGGCCTTCAGTGGGTGGGATGGATCAATCCTAAGACAGGACAGCCAAACAATCCTCGTCAATTTCAAGGTAGAGTCAGTCTGACTCGACACGCGTCGTGGGACTTTGACACATTTTCCTTTTACATGGACCTGAAGGCACTAAGATCGGACGACACGGCCGTTTATTTCTGTGCGCGACAGCGCAGCGACTATTGGGATTTCGACGTCTGGGGCAGTGGAACCCAGGTCACTGTCTCGTCAGGTGAGTCCTCGAGG was integrated at the XhoI restriction site.

For the promotor-less donors, we amplified from C57/BL6 genomic DNA the IgM splice acceptor using primers <u>CGATGCATAGGGACAAACGTGTAGAGGGATCTCCTGTCTGACAG</u> (forward) <u>and TTCGCGCGCATTTGGGAAGGACTGACTCTCTGAG</u> (reverse) <u>and replaced the PolyA-ED promoter cassette with the resulting product directly followed by a Furin-GSG-P2A cassette. For the single chain donors, we replaced the Furin-GSG-P2A between the two chains coding sequences with a variant of a previously described sequence^2^: GGCGCTGGCAGCGGCGCTTCCTGGAGTCACCCTCAGTTCGAGAAAGGCGCCTCTGGTGGATCTGGCGGCGCATCCTGGAGCCATCCACAATTTGAAAAAGGCGCTAGCGGAGGCTCTGGCGGAAGTGGCGGAGCTTCTTGGAGCCATCCGCAGTTCGAGAAAGGTGCTAGCGGCGGC</u>

### rAAV production

rAAV-DJ and rAAV-6 donors were produced in 293t cells by transient transfection. In short, 10-14 15cm dishes were transfected when cells were 80% confluent pAd5 (helper plasmid), rAAVDJ or rAAV6 genome plasmid and Donor plasmid at a 3:1:1 ratio in Polyethylenimine (PEI)(Merck). In total each plate was teransfected with 41,250ng of DNA. Purification was performed with the AAVpro Extraction Kit (Takara) according to manufacturer protocol and titer quantification by qPCR with SYBRGreen (ThermoFisher).

### Cell lines cultures

For cell lines, ImProB^3^, A-20^4^ and i.29 IgA^5^ electroporations were performed with 18.3pmol Cas9 and 22pmol gRNA, at 1.0E5cells/μl in OptiMEM using 10μl tips in a Neon electroporation system (Invitrogen). For the human cell line, parameters were: 1350v 30ms 1pulse and for the murine cell lines: 1600v 20ms, 1pulse. All cell lines were grown in 1640 RPMI supplemented with 10% HI FBS, 50μM β-Mercaptoethanol and P/S. Transductions were performed with a 50,000 MOI of rAAV-DJ for murine cell lines, and a 130,000 MOI of rAAV-DJ for human cell line. Efficiency of editing was determined 3 days following electroporation.

For plasmid electroporations we used 3μg plasmid DNA/1E06 cells/10μl Neon tip.

### Primary human cultures

For human cells, whole blood was obtained from the Israeli blood bank (Magen David Adom, Sheiba Medical Center) in accordance with Tel Aviv University Review Board, and PBMCs were extracted using Lymphocyte Separation Medium (mpbio). Remnants of red blood cells were lysed as described above. B cells were enriched using the negative selection Easysep human B cell isolation kit (Stemcell) and plated in 1640 RPMI supplemented with 10% HI FBS, 50μM β-Mercaptoethanol, P/S, 2μg/ml RP105 (LEAF, anti-human CD180, Biolegend) and 10ng/ml IL7.

For human primary cells, electroporation parameters were: 1750v 20ms 1pulse in a Neon electroporation system (Invitrogen) at 4.0E5 cells/μl in buffer R using 10μl tips. For RNP, Cas9 (IDT) and gRNA (IDT): complexes assemblies were generated 20min prior to transfection with 18.3pmol Cas9 and 66pmol gRNA per 1E06 cells. Transductions were performed no later than 5min following electroporation with a 10,000 MOI of rAAV-6. Efficiency of editing was determined 2 days following electroporation.

### Primary murine cultures

For murine cells, whole spleens were extracted from 6 to 10 weeks old female mice and were mechanistically crushed in PBS to be filtered in a 70μm Cell Strainer (Corning). Following Red Blood Cell lysis (Biolegend), cells were plated at 3.0E6 cells/ml in 1640 RPMI supplemented with 10% HI FBS, 50μM β-Mercaptoethanol, P/S, 10μg/ml LPS (SantaCruz Biotechnology) and 10ng/ml IL4 (Peprotech) was added immediately after extraction.

Cells were cultured 16-24h and washed in PBS before transfections and plated in the same activation medium without P/S for 8-16h following electroporations. Parameters were 1350v 30ms 1pulse at 4.0E5 cells/μl in buffer R for 10μl tips. For RNP: Cas9 (IDT) and gRNA (IDT) complexes assemblies were generated 20min prior of transfection with 18.3pmol Cas9 and 38pmol gRNA per 1E06 cells. Transductions were performed no later than 5min following electroporation with a 10,000 MOI of rAAV-6 and cells were analyzed 2 days following electroporation. For co-culture with CD40LB feeders, splenic lymphocytes were seeded as previously described^6^.

### Flow cytometry

For samples from recipient (CD45.2) mice, cells were stained immediately following extraction. Samples were washed and resuspended in Staining Buffer (Biolegend). Antibodies were added and incubated for 20min in the dark at room temperature for binding. Samples were then washed and resuspended in Staining Buffer before reading in an Attune NxT (Life technologies) flow cytometer. For indirect Flow cytometry: samples were incubated for 5min with 1ug of the YU2.DG gp120 antigen in 100μl, washed and conjugated antibodies were added in 100μl for 15min followed by additional washing and acquisition.

For assessing ERK phosphorylation, cells were incubated for 3min at 37°C in 100μl PBS before supplementing with 5μg of the YU2.DG gp120 antigen, incubated for one more minute, immediately put on ice and diluted by a factor of 10 with ice-cold 1:1 MetOH:Acetone and incubated for 20min at −20C. Cells were then washed and stained as described above.

For viability staining, Propidium Iodide (eBioscience) was added immediately before acquisition.

For assessment of IgK ablation, we used the following equation:

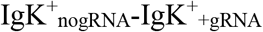

ERK phosphorylation was assessed as previously described^7^. In short, cells were washed and resuspended in PBS at 2E06 cells/ml. Cells were incubated for 3min at 37°C before supplementing by 5μg of the YU2.DG gp120 antigen, incubated for one more minute and immediately put on ice and diluted by a factor of 2 with MetOH and incubated for 15min at −20°C before staining.

Pre-gating for all experiments can be found in Fig.S15.

Data was compiled and analyzed using Kaluza Analysis 2.1 (Beckman Coulter).

Antibodies used:

FITC labeled anti-mouse CD45.1 (Invitrogen),
BV421 labeled anti-mouse IgG1 (BD Bioscience),
BV786 labeled anti-mouse IgA (BD Bioscience),
BV421 or FITC labeled anti-mouse CD19 (Biolegend),
Alexa Fluor 700 labeled anti-CD45RA/B220 (Biolegend),
APC labeled anti-mouse CD138 (Biolegend),
Pacific blue or PerCPCy5.5 labeled anti-mouse GL7 antigen (Biolegend),
PE labeled anti-mouse CD80 (Biolegend),
PE labeled anti Mo/Hu pERK1/2 (eBioscience),
Alexa Fluor 700 labeled anti mouse CD38 (eBioscience),
Alexa fluor 488 or Alexa fluor 594 or FITC or Alexa fluor 648 labeled anti-His-tag (MBL or Biolegend),
Vioblue anti-mouse IgK (MACS),
BV421 anti-human IgK (Biolegend)
PE labeled anti-human IgG Fc (eBioscience)
Anti-3BNC117 (produced in-house)
HRP conjugated anti-mouse IgM (Jackson Immunoresearch)
HRP conjugated anti-mouse IgG (Jackson Immunoresearch)
HRP conjugated anti-mouse IgG1 (Jackson Immunoresearch)
HRP conjugated anti-mouse IgA (SouthernBiotech)
HRP conjugated anti-mouse IgG2b (BioRad)
HRP conjugated anti-mouse IgG2c (BioRad)

### Enzyme Linked Immunosorbent assay

High binding micro-plates (greiner bio-one) were coated with 5μg/ml of an anti-idiotipic antibody against 3BNC117 or 2μg/ml of the YU2.DG gp120 antigen in PBS overnight at 4°C. Plates were washed with PBST and blocked for two hours with 5% BSA in PBST and washed again. Plates were then applied with detection antibodies, anti-mouse IgA(abcam) or anti-mouse IgG or anti-mouse IgG1 or anti-mouse IgM (Jackson ImmunoResearch) at 2μg/ml in PBST and were incubated for two hours. Before detection with QuantaBlu (ThermoFisher) according to manufacturer protocols, plates were washed for an additional round. Detection was done in a Synergy M1 Plate reader (Biotek). When absolute quantitation is presented, the concentration of 3BNC117 was determined by reference to the dilution factor of the standard curve. gp120 Env proteins from THRO4156 clone 18 SVPB15 (GenBank accession number AY835448) were cloned in pcDNA3.1. The YU2.DG gp120 vector was previously described^8^ (for expression under the human CD5 leader sequence, and with a His-tag (6×) on the C terminus. Plasmids encoding gp120 were transfected into Expi293F cells at a density of 2E6 cells/ml in Expi293 Expression Medium (ThermoFisher) using ExpiFectamine (ThermoFisher) according to manufacturer protocols. Supernatants were collected 7 days later and bound with Ni-NTA Agarose (Qiagen) in 20mM Sodium Phosphate, 0.5M NaCl, 10mM Imidazole (ID). Beads were washed twice with the same buffer before mounting on gravity-flow Polypropylene columns (Biorad). Chromatography elution was performed in three fractions: 50, 100 and 200mM ID. Elutes were buffer exchanged to PBS using Amicon Ultra-15 Centrifugal Filter Units (Merck) following filtration in 0.22μm filter unit (Millex).

For 3BNC117 and anti-idiotypic antibodies, 7.5μg of a plasmid encoding the light chain and 22.5μg of plasmid encoding heavy chain were co-transfected into Expi293F cells at a density of 2.0E6 cells/ml in Expi293 Expression Medium (ThermoFisher) using ExpiFectamine (ThermoFisher) according to manufacturer protocols. Purification of the supernatant, 7 days post transfection, was performed using MabSelect (GE Healthcare) following manufacturer recommendations.

### Mouse experiments

All mouse experiments were done with approval of TAU ethical committees.

Engineered splenic B cells were transferred to 6-9 weeks old female CD45.2 C57BL/6 mice (Envigo) by retro-orbital injections at 1.5M-2.2M cells/mouse in 100 μl Mg^+2^ and Ca^+2^ supplemented PBS with 5% Horse Serum. Mice were anesthetized with 0.1mg/g Ketamine and 0.01mg/g Xylazine prior to transfer. Number of cells to be transferred was set to be 112,500 gp120 binding cells and the ratio of cells was determined by flow cytometry prior to infusion. Cells were diluted in mock cells (electroporated without gRNA and then transduced) if needed and counted in a TC-20 automatic cell counter (Biorad). For immunizations, gp120 in PBS (200μg/ml for 100μl/20μg/mice) was mixed at a 1:1 ratio with Alhydrogel 2% (Invitrogen) and injected intraperitoneally. Mice receiving non-engineered cells were adoptively transferred with 1.5M cells/mouse.

Blood samples were collected by terminal bleeding in heparin. Sera from the different mice were distilled by repeated light centrifugation and collection of the supernatant until no erythrocytes were found.

### Nucleic acid manipulations

Reverse transcriptions were executed on total RNA extracted using QuickRNA microPrep kit (Zymo Research) from 1-2E06 cells 3 days following transfection with CRISPR-RNPs and AAV transduction. Cells transfected without the gRNA, but otherwise similarly treated, were used as control. 500-1500ng of extracted RNA were used for each RT reaction using either M-MLV (Promega) or RevertAid (ThermoFisher) reverse transcriptase with oligo dT according to respective manufacturer instructions. Subsequent to RT reactions, PCR reactions were done using Maxima HotStart GreenMix at 35cycles using the following primers: CGCGCGAAACGCGGAAG (forward) and for mouse IgG2a: CCACCACAGAGGAGAAGATCCAC (reverse), for mouse IgM: CGTGGTGGGACGAACACATTTAC (reverse) for mouse IgA: CATGTGAGGCTGGCATCTGAAC (reverse).

For assessment of gRNA activity, genomic DNA was extracted using Quick DNA miniprep kit (Zymo Research) and 300-500ng genomic DNA was amplified by PCR using PrimeSTAR MAX (Takara) for 30-35cyles using the following primers: mouse IgH: GGATATTTGTCCCTGAGGGAGCC (forward) GCCATCTTGACTCCAACTCAACATTG (reverse), mouse IgK: AGTCCAACTGTTCAGGACGCC (forward) GTGTGGCTAAAAATTGTCCCATGTGG (reverse), human IgH: GCTGAGGAATGTGTCTCAGGAGC (forward) CCTCAATTCCAGACACATATCACTCATGG (reverse), human IgK: GCTGGAACAGTCAGAAGGTGGAG (forward) GCTGTCCTTGCTGTCCTGCT (reverse). Amplicon DNA was denatured and reannealed in a thermocycler prior to cleaving by T7 Endonuclease 1 (New England Biolabs) at 37°C in a 30min reaction. Proteinase K was supplemented to the reaction and incubated for an additional 15min at 37°C. Cleavage was analyzed by agarose gel electrophoresis and quantified using Biovision (Vilber Lourmat) using a rolling ball for background subtraction. Efficiency was calculated as

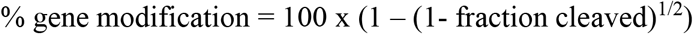

For assessment of HDR mediated integration, PCR was performed using primers CGCGCGAAACGCGGAAG (forward) ATATCACGCGTGTACACTAGCCAGTTTCGGCTGAATCCTCA (reverse) on genomic DNA extracted 2 days following transduction. Agarose (Hy-labs) was supplemented to 40mM Tris, 20mM Acetate, 1mM EDTA for a final concentration of 1-2%. Gels were run at 160v for 20-30mins. DNA ladders used were either 100bp DNA Ladder H3 RTU or 1kb DNA Ladder RTU (GeneDireX).

### Illumina sequencing and analysis

Whole spleens were extracted from the mice, mechanistically crushed in PBS and subsequently filtered in a 70μm Cell Strainer (Corning). The preparation underwent Red Blood Cell lysis (Biolegend). cDNA was generated from Quick RNA Microprep (Zymo) extracted RNA of total splenic lymphocyte populations with the RevertAid reverse transcriptase (ThermoFisher) using Oligo-dT primers. For Initial PCR amplification, VH fragments were amplified with the proofreading PrimeStarMAX polymerase (Takara) for 40 cycles using primers: TCGTCGGCAGCGTC AGATGTGTATAAGAGACAG**X** CAGGTTCAGCTGCTGCAATCTG (forward) and GTCTCGTGGGCTCGGAGATGTGTATAAGAGACAG**X** CTGACGAAACAGTAACTTGGGTTCC, where X stands for 0-7 degenerate positions (“N”s). Amplicons were purified using AMPureXP beads (Beckman Coulter) at a 0.7:1 ratio. The subsequent 8 cycles PCR reaction was performed with the PrimeStarMax polymerase (Takara) using primers: CAAGCAGAAGACGGCATACGAGAT**X** GTCTCGTGGGCTCGG and AATGATACGGCGACCACCGAGATCTACACTAGATCGCTCGTCGGCAGCGTC where X stands for i7 indexes. Libraries were purified using AMPureXP beads at a 0.7:1 ratio. Combined libraries were loaded at 5pM with 25% PhiX control (Illumina). Sequencing was performed in a high-throughput MiSeq machine using either a v2 Nano reagent kit 2×250bp or a v3 reagent kit 2×300bp (Illumina) at the Genomic Research Unit (GRU) at the Faculty of Life Sciences, Tel Aviv University. Raw fastq files were submitted to Fast Length Adjustment of Short Reads (FLASH)^9^ and resultant paired-end fasta files were submitted to High V-Quest for V gene sequence alignment and germ-line gene assignment provided by the international ImMunoGeneTics database (IMGT) ^10^. IMGT aligned sequences were then processed by a series of quality control filters. For amino acid analyses the filters included: removal of truncated sequences, sequences containing stop codons and sequences with a similarity of less than 95% with the original 3BNC117 sequence. For both nucleic acid and amino acid sequences, we trimmed the 5’ and 3’ ends to reduce sequencing bias (Fig.S13F). For nucleic acid analyses, we kept non-productive sequences but removed high InDel count sequences. For polarity analyses, we summed frequency of the top 10 clones among mutant sequences. For the analysis of synonymous vs non-synonymous mutations, we used the biopython^11^ alignment implementation of the Needleman-Wunsch algorithm to pairwise align the mutant V_H_ sequences with the V_H_ of 3BNC117-opt sequence.

For the N-Syn/Syn ratio, we used the following equation: 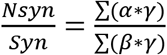, where α is the number of non-synonymous mutations in a sequence, γ is the frequency of that sequence and β is the number of synonymous mutations in that sequence.

Clustal Omega^12^ was used for tree constructions (Fig. S14A). Alignment for sequences was performed via SnapGene v5.0.7.

### Statistical Analysis

Statistical Analysis was performed using GraphPad Prism 8 to calculate p-values with two- or three-way ANOVA or two tailed Student’s t-tests. One sample t-test was performed by setting expected value as for the *in-vitro* controls. For ANOVA, Tukey’s or Dunnett’s multiple comparisons were performed as indicated.

## List of Supplementary Materials

Figures S1-S15

